# Affibody-conjugated hyaluronic acid hydrogels within collagen sponges enhance dose-dependent BMP-2-mediated bone formation and repair of critically sized femoral defects

**DOI:** 10.64898/2026.11.19.767909

**Authors:** Yan Carlos Pacheco, Andy J. Huang, Nick J. Willett, Marian H. Hettiaratchi

## Abstract

1.

Bone morphogenetic protein-2 (BMP-2) is a potent osteoinductive growth factor; however, its burst release from clinical collagen sponge carriers can cause inflammation and ectopic bone formation. Alternative delivery vehicles slow BMP-2 protein release through increased crosslinking density, which stiffens the polymer network and can restrict cell infiltration required for bone formation. Here, we introduce a BMP-2 delivery system in which a hyaluronic acid (HA) hydrogel containing BMP-2-specific affibodies is formed within clinically used collagen sponges to retain BMP-2 through protein-material affinity interactions rather than physical entrapment. We developed a maleimide functionalized, hydrazone crosslinked HA hydrogel such that affibody conjugation is chemically orthogonal to network formation, allowing affibody density to be varied independently of crosslinking density. Forming this hydrogel within a collagen sponge produced composite scaffolds with improved surgical handling and BMP-2 retention. Moreover, *in vitro*, composite scaffolds exhibited minimal swelling while retaining more BMP-2 than the collagen sponge alone. *In vivo*, composite scaffolds conjugated with BMP-2-specific affibodies improved ectopic mineralization in a BMP-2 dose-dependent manner. In a critically sized femoral defect, composite scaffolds with and without affibodies significantly increased bone volume relative to the collagen sponge. Thus, this composite scaffold effectively integrates the clinically used collagen sponge with a tunable HA hydrogel and BMP-2-specific affibodies to improve BMP-2-mediated bone regeneration and expand the use of naturally derived biomaterials for bone healing.

**Graphical Abstract:** 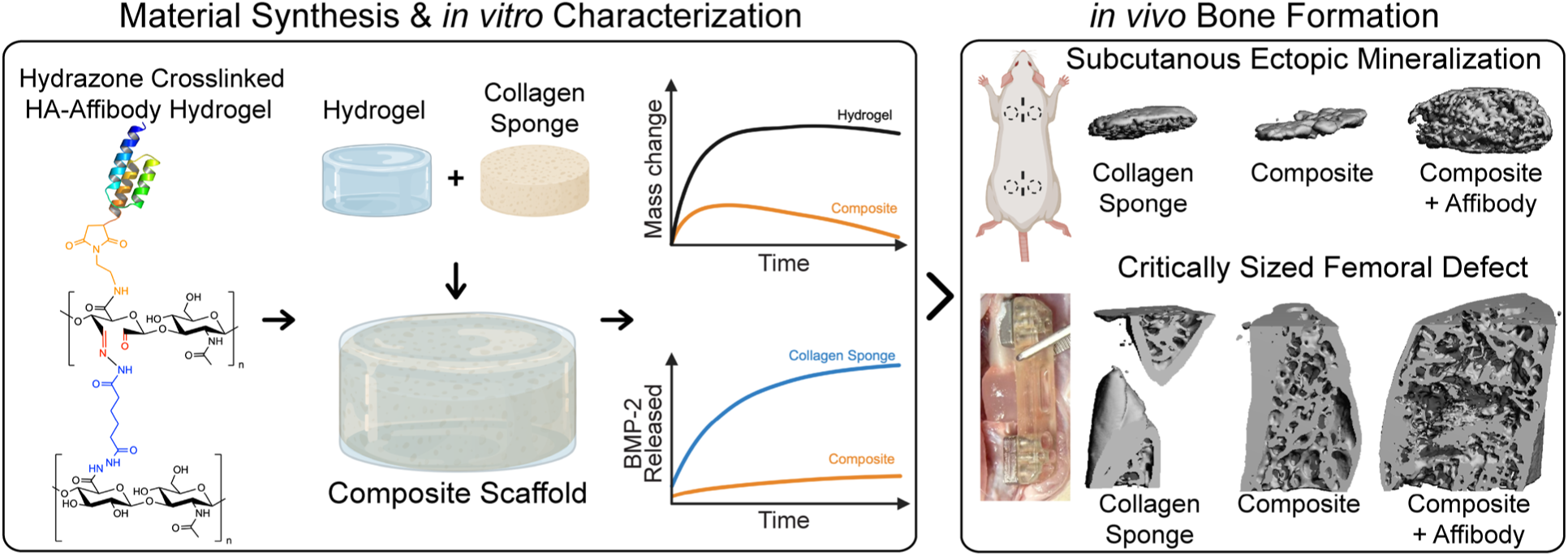

## 2. Introduction

Bone is one of the few tissues that heals without scars, yet bone injuries resulting from significant trauma frequently exceed its intrinsic healing capacity and require surgical intervention.[1] Critically sized bone defects and the estimated 120,000-450,000 spinal fusions that result in non-fusion or off-target effects each year, represent a large population of patients in which bone does not regenerate on its own.[2,3] The standard of care for long bone and spinal defects is a multi-step procedure involving bone stabilization and fixation, followed by autologous or allogeneic bone graft. Autologous bone graft remains the gold standard for treating both types of bone defects because it is simultaneously osteogenic, osteoinductive, and osteoconductive. Yet, autologous bone graft supply is limited, and graft harvest can cause donor site pain, intraoperative blood loss, infection, nerve injury, and iliac fracture.[2] Bone allografts address the supply constraint, but allografts lose osteogenic activity during sterilization and raise concerns about disease transmission and immune rejection. Ceramic substitutes are only osteoconductive and contain no osteoinductive or osteogenic components.[4] Limitations of current bone grafting strategies have driven the adoption of recombinant human bone morphogenetic protein-2 (BMP-2), a potent osteoinductive growth factor, for anterior lumbar interbody fusion. In a meta-analysis of 40 studies, adding BMP-2 to autologous bone grafts increased fusion rates from 92.5% to 96.6%, which is a modest improvement that must be weighed against the complications associated with the BMP-2 doses required to achieve it, including postoperative inflammation, ectopic bone formation, and graft subsidence.[5,6] These complications are tied to how BMP-2 delivered; through an absorbable bovine type I collagen sponge, which binds the protein through weak electrostatic interactions.[4,7,8] Because the collagen sponge retains BMP-2 poorly at the implant site, supraphysiological BMP-2 doses are required to induce osteogenesis, which can cause soft tissue inflammation, heterotopic bone formation, and other complications.[9–12]

Improving the collagen sponge for BMP-2 delivery is challenging because an effective BMP-2 carrier must satisfy two competing requirements. First, because bone regeneration proceeds by cellular invasion of the implant, the carrier must remain soft and porous enough to permit progenitor cell infiltration, vascular ingrowth, and matrix deposition. Second, the carrier must retain BMP-2 in the injury site long enough to stimulate osteogenesis. The most common way to slow protein release is to increase polymer crosslinking density or reduce biomaterial mesh size, which stiffens the network and restricts cell infiltration.[13–17] Affinity-based protein delivery addresses this issue by improving protein retention within a carrier through molecular interactions between the protein and the material itself rather than physical entrapment. Extracellular matrix molecules that naturally bind BMP-2, such as heparin, can be incorporated into biomaterials to prolong BMP-2 retention and enhance its bioactivity.[18] Incorporating heparin microparticles into alginate hydrogels improved the spatial localization of BMP-2-induced bone formation within a critically sized femoral defect in rats.[19] However, heparin binds other growth factors in addition to BMP-2, as well as numerous serum proteins that can displace bound BMP-2 and accelerate its release *in vivo*.[20] Additionally, scaffolds composed of collagen and heparin have yielded mixed results for BMP-2 delivery for bone repair, which may be attributed to the dose of heparin used and how heparin is immobilized within the material.[21] Because heparin binds promiscuously to a number of different proteins, BMP-2 retention on heparin biomaterials cannot be tuned without simultaneously changing how the carrier interacts with every other protein it binds.[20,22]

To overcome some of the drawbacks of natural affinity-based materials, our lab has engineered highly specific affinity-based protein binders. Affibodies are small (∼7 kDa) protein domains that can be engineered to have affinity towards a chosen target. Our group previously identified BMP-2-specific affibodies and demonstrated that conjugating a moderate-affinity affibody (dissociation constant, K_D_ = 10.7 nM) to a hydrogel slowed BMP-2 release while preserving BMP-2 bioactivity.[23] Affibody-conjugated polyethylene glycol maleimide (PEG-Mal) hydrogels were used to deliver BMP-2 to a femoral defect, increasing BMP-2 retention and achieving complete defect bridging over 12 weeks.[24] However, in this approach, cysteine-terminated affibodies were conjugated to maleimide functional groups, which were also used for hydrogel crosslinking. Conjugating 1 mM of cysteine-terminated cell-adhesive peptides and 0.65 mM of affibodies to the maleimides consumed 16.5% of crosslinking sites, altering hydrogel stiffness, swelling, and mesh size. Using orthogonal chemical reactions for affinity binder conjugation and hydrogel crosslinking would instead enable tuning of hydrogel physicochemical and mechanical properties, independent of affibody density. Moreover, orthogonal chemical reactions could increase the total number of affibodies that could be conjugated to the polymer, overcoming previous limitations in conjugating enough affibodies to control BMP-2 release *in vivo* without compromising hydrogel gelation.[25] Synthetic PEG hydrogels also require conjugation of cell-adhesive peptides alongside matrix metalloprotease (MMP) degradable crosslinks to support cell attachment, infiltration, and hydrogel remodeling, whereas HA is itself a natural extracellular matrix that supports cell interactions and is degraded by endogenous hyaluronidases.[16,26]

We developed an HA hydrogel with three key features: 1) a maleimide functional group for conjugation of cysteine-terminated BMP-2-specific affibodies via thiol-Michael addition, 2) hydrazone crosslinking that is chemically orthogonal to affibody conjugation, and 3) *in situ* formation of the hydrogel within the clinically used collagen sponge. These features address two key requirements for improving upon the collagen sponge delivery vehicle without replacing it entirely: retaining BMP-2 through affinity interactions and preserving the porosity needed for cell infiltration.

In this study, we investigated the efficacy of composite collagen-HA scaffolds to improve BMP-2-mediated bone repair in a critically sized femoral defect rat model. We hypothesized that BMP-2-specific affibody conjugation would increase *in vitro* BMP-2 retention and enhance *in vivo* BMP-2 dose-dependent bone formation relative to both the collagen sponge and the composite scaffolds without affibodies. We first established that maleimide functionalization and affibody conjugation left hydrazone crosslinking intact by measuring compressive modulus, swelling, degradation, and BMP-2 release from hydrogels and composite scaffolds. Having confirmed that affibody conjugation and addition of a collagen sponge did not compromise hydrogel properties, we asked whether the retention it provides translates into bone formation in a subcutaneous ectopic model with implants loaded with either 2.5 or 5.0 µg of BMP-2. Finally, we implanted composite scaffolds with and without affibodies, loaded with 1.0 µg of BMP-2, into critically sized rat femoral defects and compared their ability to repair bone with BMP-2-loaded collagen sponges. We assessed bone regeneration using x-ray radiography, micro-computed tomography, and histology. Together, these studies establish a novel composite scaffold that combines clinically relevant collagen sponges with novel BMP-2-specific affinity binders to improve bone regeneration in large bone defects.

## 3. Materials and Methods

### 3.1. Materials

Sodium hyaluronate (100 kDa) was purchased from LifeCore Biomedical LLC (Chaska, MN). Adipic acid dihydrazide (ADH) was purchased from Spectrum Chemical (Gardena, CA). 1-Hydroxybenzotriazole hydrate (HOBt) was purchased from Chem-Impex (Wood Dale, IL). 1-Ethyl-3-(3-dimethylaminopropyl)carbodiimide (EDC) was purchased from G Biosciences (St. Louis, MO). N-(2-aminoethyl)maleimide HCl and 4-(4,6-dimethoxy-1,3,5-triazin-2-yl)-4-methylmorpholinium chloride (DMTMM) were purchased from AmBeed (Buffalo Grove, IL). Sodium periodate was purchased from Sigma-Aldrich (St. Louis, MO). Ellman’s reagent (5,5’-dithio-bis-[2-nitrobenzoic acid]) was purchased from Fisher Scientific (Waltham, MA). Recombinant human bone morphogenetic protein-2 (BMP-2) and BMP-2 DuoSet ELISA were purchased from R&D Systems (Minneapolis, MN).

### 3.2 Synthesis of HA adipic acid dihydrazide (HA-ADH)

1% (w/v) sodium hyaluronate (0.125 g, 0.000330 mol) was dissolved in double-distilled water (ddH_2_O) and stirred until completely dissolved. 10 molar equivalents of adipic acid dihydrazide (ADH) (0.575 g, 0.00330 mol) were added to the reaction and stirred until dissolved. One molar equivalent of 1-hydroxybenzotriazole (HOBt) (0.0446 g, 0.000330 mol) was added, and the pH was adjusted to 4.75 using 0.1 M NaOH/HCl. One molar equivalent of 1-Ethyl-3-(3-dimethylaminopropyl) carbodiimide (EDC) (0.0632 g, 0.000330 mol) was added to the solution. The pH was titrated back to 4.75 and monitored for 2-4 h until stable. The reaction was then stirred for 2 days before being dialyzed against 0.1 M sodium chloride for 2 days (six complete water changes), followed by ddH_2_O for an additional 2 days (six complete water changes). The solution was passed through a 0.22 µm syringe filter, frozen at −80 °C, and lyophilized.

### 3.3 Synthesis of HA maleimide (HA-Mal)

1% (w/v) sodium hyaluronate (0.500 g, 0.001319 mol) was dissolved in ddH_2_O and stirred until completely dissolved. 3 molar equivalents of DMTMM (1.095 g, 0.003958 mol), dissolved in 3 mL of ddH_2_O, were added dropwise to the reaction and allowed to dissolve completely between additions. 2 molar equivalents of N-(2-aminoethyl)maleimide HCl (Mal-NH_2_) (0.466 g, 0.002639 mol) were similarly added to the reaction. The reaction was then stirred overnight, dialyzed against 0.1 M sodium chloride for 2 days (six complete water changes), and then dialyzed against ddH_2_O for an additional 2 days (six complete water changes). The solution was passed through a 0.22 μm syringe filter, frozen at −80 °C, and lyophilized.

### 3.4 Oxidation of HA maleimide (HA-Mal-Ox)

HA-Mal was oxidized to reveal aldehyde groups using methods previously described for oxidizing sodium hyaluronate.[25] 2% (w/v) HA-Mal (0.500 g, 0.001319 mol) was dissolved in ddH_2_O in a round-bottom flask and stirred until completely dissolved. The flask was covered with foil, and 0.6 molar equivalents of sodium periodate (0.169 g, 0.0007914 mol) were added. The reaction was stirred for 4 h in the dark, quenched with 5 molar equivalents of propylene glycol (VWR Chemicals, Radnor, PA) (0.502 g, 0.006596 mol), and stirred for an additional 1 h. The solution was then dialyzed against 0.1 M sodium chloride for 2 days (six complete water changes), followed by ddH_2_O for an additional 2 days (six complete water changes). The solution was passed through a 0.22 μm syringe filter, frozen at −80 °C, and lyophilized.

### 3.5 Determination of polymer degree of modification (DOM)

Degree of modification of the HA polymer was determined as previously described.[25,27] The DOM for HA-ADH and HA-Mal was quantified via proton nuclear magnetic resonance spectroscopy (^1^H-NMR, 500 MHz, Bruker, USA).[25] The DOM for HA-ADH was calculated by integrating the aliphatic chain peaks from 2.5 to 2.1 ppm and 1.7 to 1.44 ppm (8H) and normalizing to the N-acetyl methyl group peaks on HA from 1.8 to 2 ppm (3H).[27] The DOM for HA-Mal was calculated by integrating the vinyl peaks of the maleimide functional groups from 6.8 to 6.9 ppm (2H) and normalizing to the N-acetyl methyl group peaks on HA.[28,29] Representative ^1^H-NMR spectra for HA-ADH and HA-Mal are provided in the supplemental information (Fig. S1-2).

To determine the DOM for oxidized polymers, we performed a hydroxylamine hydrochloride titration by dissolving 50 mg of oxidized polymer in 10 mL of 0.25 N hydroxylamine hydrochloride containing 0.05% (w/v) methyl orange reagent for 2 h.[25,30,31] The pH of the solution was monitored and titrated with 0.1 M NaOH until it reached the pH of the stock 0.25 N hydroxylamine solution. Equation 1 was used to calculate the DOM, or the percentage of HA disaccharide units that contained aldehydes.

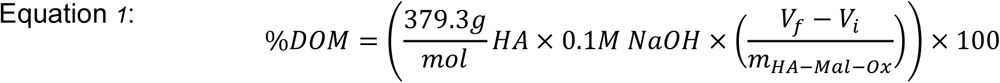

where 379.3 g/mol is the molecular weight of one HA disaccharide unit, 0.1 M is the molarity of NaOH used for titration, V_f_ and V_i_ are the final and initial titration volumes in liters, and m_HA-Mal-Ox_ is the mass (g) of HA-Mal-Ox initially dissolved for titration.

### 3.6 Solid phase affibody synthesis

We synthesized the BMP-2-specific affibody using the Liberty Blue 2.0 microwave peptide synthesizer (CEM Corporation), as previously described.[24,32] Briefly, we modified the affibody sequence with a penultimate cysteine and a C-terminal glycine and synthesized it on Fmoc-Gly-Wang resin. Following synthesis, the crude affibody was cleaved from the resin, precipitated using cold diethyl ether, and resolubilized with a 10% (v/v) acetic acid solution. The affibody was then lyophilized and dissolved in 3% (v/v) acetonitrile (with 0.1% (v/v) trifluoroacetic acid) at 10 mg/mL. The dissolved affibody solution was purified using a Prodigy high-performance liquid chromatography (HPLC) system (CEM Corporation) by running a 3-70% gradient of acetonitrile in H_2_O with 0.1% (v/v) trifluoroacetic acid. Affibodies were lyophilized and stored at −20 °C until use. We confirmed the size and purity of the affibody using matrix-assisted laser desorption/ionization time-of-flight (MALDI-TOF) mass spectrometry (Bruker Smart LS). Representative MALDI-TOF spectra of crude and purified affibody are provided in the supplemental information (Fig. S3).

### 3.7 HA-Mal hydrazone hydrogel formation

HA-Mal-Ox and HA-ADH polymers were reconstituted at 2% (w/v) in pH 7.4 phosphate-buffered saline (PBS). 100 µL hydrogels were prepared by mixing 50 µL of each polymer in an 8 mm diameter cylindrical silicone mold and allowing them to crosslink for 10 minutes. For hydrogels containing affibodies, HA-Mal-Ox polymers were dissolved at 2.5-3% (w/v) and mixed with affibodies to achieve a final HA-Mal-Ox concentration of 2% (w/v). Similar to HA hydrogels without affibodies, 50 µL of HA-Mal-Ox polymer solution containing affibodies was mixed with 50 µL of HA-ADH polymer solution in 8 mm diameter molds to create 100 µL hydrogels. Ellman’s reagent was used to detect free thiols and quantify affibody concentration before and after mixing with HA-Mal-Ox polymers as previously described.[27]

### 3.8 Compressive moduli of HA-Mal hydrogels

We evaluated the compressive moduli of the hydrogels using a DHR-30 hybrid rheometer (TA Instruments). 100 µL 2% (w/v) HA hydrogels were made with HA-ADH and either HA-Ox or HA-Mal-Ox in 8 mm molds, as described above. For affibody-conjugated hydrogels, BMP-2-specific affibodies were added to HA-Mal-Ox solution before hydrogel formation. The hydrogels were then allowed to crosslink overnight at 4 °C in a humid chamber before being removed from the molds and placed on the rheometer. Hydrogels were compressed within 8 mm parallel plates to 50% of their original height at room temperature, and the slope of the linear portion of the stress-strain curve was used to calculate the compressive modulus. A representative stress-strain curve of an HA-Ox hydrogel is provided in the supplemental information (Fig. S4).

### 3.9 Hydrogel and composite scaffold mass change and BMP-2 release

The swelling, degradation, and BMP-2 release of hydrogels and composite scaffolds were assessed over 21 days. Five construct groups were prepared (n = 4 per group): 1) HA hydrogels, 2) Affibody-conjugated HA hydrogels, 3) collagen sponges (CS), 4) CS + HA hydrogels, 5) CS + affibody-conjugated HA hydrogels. For affibody-conjugated HA hydrogels, 1.92 nmol of BMP-2-specific affibodies were conjugated for both mass change and BMP-2 release experiments.

For mass change assessment, hydrogel constructs were crosslinked overnight at 4 °C in a humid chamber, then removed from the mold and weighed to record an initial mass. For collagen sponge constructs, 8 mm cylindrical disks were cut from a sheet of Medtronic bovine type I collagen sponge using an 8 mm biopsy punch and placed into cylindrical molds. Collagen sponges were hydrated with 100 µL of PBS and weighed to record an initial mass. For composite scaffolds with and without affibodies (CS + HA hydrogels, CS + affibody-conjugated HA hydrogels), 8 mm collagen sponge disks were placed into cylindrical molds, and 50 µL of each polymer solution were pipetted directly onto the sponge, allowing the hydrogel to crosslink within the collagen sponge matrix. Composite scaffolds were then weighed to record an initial mass. All constructs were incubated in 1 mL of PBS at 37°C. After 1, 3, 5, 7, 14, and 21 days, PBS was removed, excess liquid was gently blotted from each construct using a Kimwipe™, and the constructs were weighed to record their wet masses. Constructs were transferred to fresh PBS following each measurement. One HA hydrogel was lost during handing, leaving n = 3 for that group.

For BMP-2 release experiments, 100 ng of BMP-2 were mixed into a 2% (w/v) solution of HA-ADH polymer prior to hydrogel crosslinking for 30 minutes at room temperature in silicone molds. Collagen sponges were hydrated with 100 µL of PBS containing 100 ng of BMP-2. For composite scaffolds, 8 mm collagen sponge disks were placed into molds, and 50 µL of 2% (w/v) HA-ADH with 100 ng of BMP-2 and 50 µL of 2% (w/v) HA-Mal-Ox were pipetted directly onto the sponge. After crosslinking and BMP-2 loading, constructs were transferred to individual 2 mL microcentrifuge tubes. 400 µL of 0.1% (w/v) bovine serum albumin (BSA) in PBS were added to each tube. After 10 minutes, 200 µL of supernatant were collected to calculate BMP-2 encapsulation. Then, 700 µL of fresh 0.1% (w/v) BSA in PBS were added to each tube, and the constructs were incubated at 37°C. At days 0, 1, 3, 5, 7, 14, and 21, 200 µL of supernatant were collected and replenished with an equal volume of fresh 0.1% (w/v) BSA in PBS. BMP-2 concentrations in collected samples were quantified using a BMP-2-specific ELISA per the manufacturer’s instructions. BMP-2 encapsulation efficiency was calculated by comparing the amount recovered in the wash to the amount of BMP-2 added.

### 3.10 Subcutaneous implantation of composite scaffolds for bone mineralization

Collagen sponges with and without HA hydrogels were implanted in the dorsal subcutaneous space of 6-7 week old female Sprague Dawley rats (Charles River Laboratories) as previously described to investigate the effect of BMP-2 dose on ectopic mineralization *in vivo*.[19,24] All animal procedures were approved by and conducted in accordance with the University of Oregon’s Institutional Animal Care and Use Committee (IACUC). All reagents were sterile-filtered through a 0.22 µm filter and handled in a biosafety cabinet using aseptic techniques. Silicone molds were cleaned with soapy water followed by a 1 h soak in 70% (v/v) ethanol in ddH_2_O. Three different constructs were prepared and loaded with either 2.5 µg or 5.0 µg of BMP-2 for a total of six groups (n = 6 per group): 1) 2.5 µg BMP-2 + collagen sponge (CS), 2) 2.5 µg BMP-2 + CS + HA hydrogels, 3) 2.5 µg BMP-2 + CS + affibody-conjugated HA hydrogels, 4) 5.0 µg BMP-2 + collagen sponge (CS), 5) 5.0 µg BMP-2 + CS + HA hydrogels, 6) 5.0 µg BMP-2 + CS + affibody-conjugated HA hydrogels. For affibody-conjugated HA hydrogels, HA-Mal-Ox polymer was mixed with a 500-fold molar excess of affibodies relative to the BMP-2 dose (48 nmol and 96 nmol of affibodies for 2.5 µg and 5.0 µg of BMP-2, respectively). Collagen sponge + HA hydrogels with and without affibodies were fabricated and loaded with BMP-2 similar to BMP-2 loading for BMP-2 release experiments. The final volume of all constructs was 100 µL.

The fur on the dorsal aspect of the rats was shaved, and the surgical site was aseptically prepared using 70% (v/v) isopropyl alcohol and 4% (v/v) chlorhexidine solution. The rats were anesthetized using inhalant isoflurane anesthesia. Two 1.5 cm incisions were made along the dorsum of each rat: one midline between the shoulder blades and the other just cranial to the pelvis. Blunt dissection was used to create two subcutaneous pockets per incision. Constructs were implanted in the subcutaneous pockets, and the incisions were closed with wound clips. Treatment groups were assigned to implant locations in a randomized order, and the surgeon was blinded to construct identity during implantation.

After 28 days, the rats were euthanized via carbon dioxide asphyxiation following the recommendations of the American Veterinary Medical Association (AVMA) Guidelines for the Euthanasia of Animals. The constructs were explanted from the subcutaneous space and transferred to a 24-well plate. X-ray radiographs were taken of each explanted construct (Faxitron MX20, 40 kV, 7 s). Constructs were then fixed in 10% (v/v) neutral buffered formalin in PBS for 24 h at 4°C and then rinsed with PBS three times. Construct mineral volume was determined using micro-computed tomography (µCT) (VivaCT80, Scanco Medical) with a 31.2 µm voxel size at 70 kVp voltage, 114 µA current, and 375 ms integration time.

### 3.11 Cryosection histology of subcutaneous implants

Representative samples from each group were selected for histological processing based on the acquired µCT data. After µCT analysis, samples were decalcified in Cal-Ex^™^ (Fisher Scientific) overnight at room temperature with slight agitation. Radiographs were taken to confirm no mineral remained, and samples were rinsed with PBS three times and submerged in 15% (w/v) sucrose in PBS at room temperature with slight agitation overnight. The solution was then changed to 30% (w/v) sucrose in PBS and incubated at 4°C overnight. Constructs were embedded in optimal cutting temperature (OCT) compound (Fisher Scientific) and sectioned into 10 µm sections using a cryostat (Leica Biosystems, CM 3050). Sections were placed on glass slides and stored at −20°C until staining.

Sample slides were allowed to reach room temperature before Hematoxylin & Eosin (H&E) staining for gross tissue morphology. Briefly, slides were submerged in PBS for three changes, rinsed in distilled water, and stained with Hematoxylin for 3 minutes. Slides were rinsed multiple times with tap water, dipped in acid alcohol, and rinsed again before being stained with Eosin-Y for 1 minute. Slides were then dehydrated through a serial ethanol ladder and cleared in xylene before a coverslip was added. Slides were imaged using a 5x objective with brightfield scanning mode (Leica Thunder). Multiple images for each sample were stitched together using the Leica LAS X automated multiple tile capture feature.

### 3.12 Femoral bone defect surgery

Composite scaffolds made with HA hydrogels with and without BMP-2 affibodies were implanted in critically sized femoral bone defects in rats to assess BMP-2-induced bone repair compared to clinically used collagen sponges. Three different constructs were prepared and loaded with 1.0 µg of BMP-2 (n = 4-6 per group): 1) collagen sponges (CS), 2) CS + HA hydrogels, 3) CS + affibody-conjugated HA hydrogels. Silicone molds (5 mm diameter, 7.5 mm height) were fabricated and sterilized to produce composite scaffolds that could be press-fit into the defect space. For composite CS + HA hydrogels, two 5 mm collagen sponges were cut and placed in molds, and 75 µL of each polymer solution were added as described above. For affibody-conjugated HA hydrogels, HA-Mal-Ox polymer was mixed with a 500-fold molar excess of affibodies (19.2 nmol) relative to the 1.0 µg BMP-2 dose. BMP-2 was loaded by mixing HA-ADH with BMP-2 prior to crosslinking. The final volume for all constructs was 150 µL. Collagen sponges without hydrogels were rehydrated with 150 µL of PBS solution containing 1.0 µg BMP-2.

Unilateral critically sized segmental bone defects were created in the femurs of 13-14 week-old female Sprague Dawley rats (Charles River Laboratories) as previously described.[19,24] Briefly, a skin incision was made on the lateral aspect of the hind left leg followed by blunt dissection to expose the femur. An internal polysulfone fixation plate was placed on the femur and secured in place by a C-clamp. The plate was then fixed with four screws. An oscillating surgical saw was used to create a 6-mm-wide, full-thickness mid-diaphyseal bone defect. Collagen sponges and composite scaffolds were placed within the defect region. The surrounding musculature was closed using resorbable sutures, and the skin incision was closed using wound clips. After surgery, rats were monitored for signs of distress and evaluated for ability to bear weight on the surgical limb for 7 days post-surgery and biweekly thereafter, with fixation plate failure defined as a criterion for exclusion from the study. Three animals were excluded due to fixation plate failure (two from the collagen sponge group and one from the composite collagen sponge and HA hydrogel group), resulting in final group sizes of n = 4, 5, and 6 for the collagen sponge, collagen sponge + HA hydrogel, and collagen sponge + affibody-conjugated HA hydrogel groups, respectively.

### 3.13 Radiographs and µCT

Longitudinal radiographs were taken at 4, 8, and 12 weeks after surgery (Faxitron MX20, 40 kV, 7 s) to assess defect bridging. A defect was scored as bridged when continuous mineralized tissue spanned the full length of the defect. *In vivo* µCT scans (VivaCT80) were performed 6 weeks after surgery using 48 µm voxel size, 55 kVp voltage, 145 µA current, and 650 ms integration time. *Ex vivo* µCT scans were performed using 24 µm voxel size, 55 kVp voltage, 145 µA current, and 750 ms integration time. Contours of the entire 6 mm defect region were drawn beginning at the intact bone ends to include all new bone formed. Mineral was quantified within the central 5.5 mm of the defect region contoured.

### 3.14 Paraffin embedded histology of femurs

Representative femurs from each group were selected based on the mean µCT bone volume measurement for each group at 12 weeks and fixed in 10% (v/v) neutral buffered formalin in PBS for 48 h. Femurs were then rinsed with PBS three times and decalcified with Cal-Ex™ (Fisher Scientific) for 3 days at room temperature with slight shaking and replacement of the solution on day 2. Radiographs were taken daily to monitor decalcification. After decalcification, samples were rinsed three times with PBS, followed by three rinses with ddH_2_O for 5 minutes each. Samples were then dehydrated in increasing concentrations of ethanol, cleared with xylene (Fisher Scientific), infiltrated with paraffin wax, and embedded in paraffin wax blocks. Serial coronal 10 µm sections were made using a microtome (Tissue-Tek, Sakura) and mounted onto glass slides. Slides from each femur were stained using Hematoxylin & Eosin (H&E) for gross tissue morphology, Safranin-O and Fast green (Saf-O) for proteoglycans in cartilage, and Picrosirius Red for collagen organization. Slides stained with H&E and Saf-O were imaged under the 5x brightfield scanning mode, and slides stained with Picrosirius Red were imaged at 10x magnification using polarized light (Leica Thunder). Multiple images for each sample were stitched together using the Leica LAS X automated multiple tile capture feature (LAS X).

### 3.15 Statistical analysis

All data are plotted as mean ± standard deviation. Statistically significant differences between groups for each experiment were determined by performing a one-way ANOVA or two-way ANOVA as applicable with Tukey’s post-hoc test for multiple comparisons. All datasets met the assumptions for performing an ANOVA. Mass change was analyzed using three-way repeated-measures ANOVA with collagen sponge, affibody conjugation, and time as factors. Collagen sponges alone were excluded from this analysis because they contain no HA hydrogel and would create an unbalanced three-way design. Femoral defect bone volume was compared by one-way ANOVA at 6 and 12 weeks. Radiographic bridging rates were compared using Fisher’s exact test. Because all defects treated with composite collagen sponge and HA hydrogels bridged regardless of affibody conjugation, the two composite scaffold groups were pooled (11 of 11) and compared with the collagen sponge group (2 of 4). Statistical significance is defined as p < 0.05. In figures, significance levels are indicated as *=p<0.05, **=p<0.01, ***=p<0.001, ****=p<0.0001. All analyses were performed using GraphPad Prism 11 (GraphPad Software).

## 4. Results

### 4.1 HA functionalization enables affibody conjugation and orthogonal hydrogel crosslinking

HA polymers were functionalized to enable thiol-Michael addition and hydrazone crosslinking (Fig. 1). 100 kDa HA was modified with N-(2-aminoethyl) maleimide HCl (Mal) via DMTMM-activated coupling to form HA-Mal (Fig. 1A). The percent degree of modification (DOM) for maleimide functionalization was 20.4 ± 2.6% (n=6), determined by ^1^H-NMR. HA-Mal was then oxidized to reveal aldehyde groups (HA-Mal-Ox) (Fig. 1B), and the DOM was determined to be 33.5 ± 2.9% (n=4). A separate batch of 100 kDa HA was modified with adipic acid dihydrazide (ADH) to form HA-ADH as previously described (Fig. 1C).[25] The DOM with ADH was determined to be 43.0 ± 4.0% (n=6) using ^1^H-NMR. For hydrogels conjugated with affibodies, we mixed HA-Mal-Ox polymer with BMP-2-specific affibodies at the desired concentration in PBS and allowed the reaction to proceed at room temperature. After 5 minutes, we measured the remaining thiol concentration in the affibody-polymer mixture by Ellman’s assay. No free thiols were detected, suggesting complete affibody conjugation.

**Figure 1.**
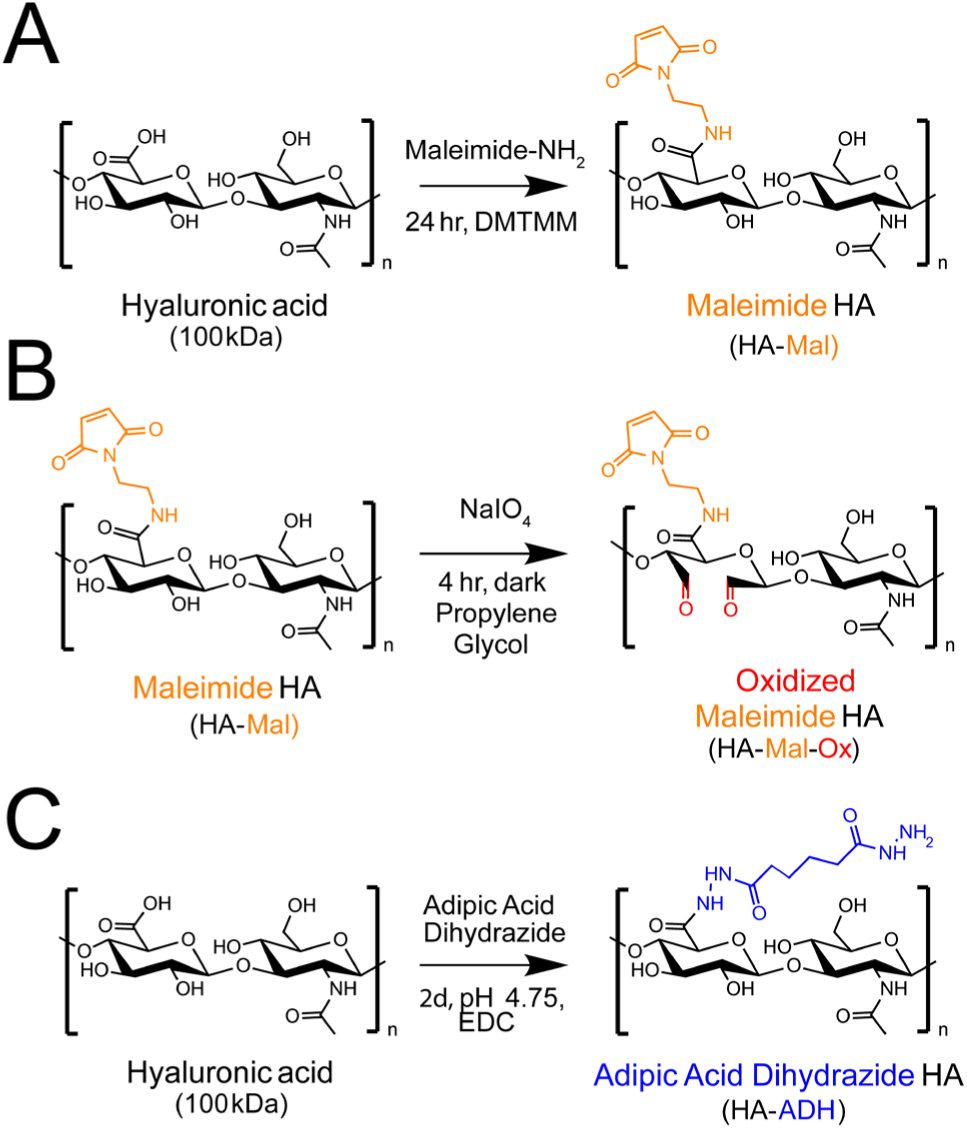
Functionalization of hyaluronic acid polymers for affibody conjugation and hydrazone crosslinking. A) Maleimide groups were introduced onto HA by coupling N-(2-aminoethyl) maleimide using DMTMM activation to form maleimide-functionalized HA (HA-Mal). B) HA-Mal polymer was subsequently oxidized with sodium periodate (NaIO_4_) in the dark for 4 h and quenched with propylene glycol to generate oxidized, maleimide-functionalized HA (HA-Mal-Ox). C) Hyaluronic acid (HA, 100 kDa) was modified with adipic acid dihydrazide (ADH) using EDC-mediated carbodiimide coupling at pH 4.75 to generate hydrazide-functionalized HA (HA-ADH).

### 4.2 Affibody conjugation does not impact HA hydrogel compressive moduli

We measured the compressive moduli of 2% (w/v) HA hydrogels without the maleimide functional group or conjugated affibodies (HA hydrogel), with the maleimide functional group (HA-Mal Hydrogel), and with two different amounts of conjugated affibodies (HA-Mal Hydrogel + 1.92 nmol Affibody, HA-Mal Hydrogel + 48 nmol Affibody) to determine whether introducing maleimides or affibodies affected hydrazone crosslinking and subsequent hydrogel stiffness. HA hydrogels displayed an average compressive modulus of 116.4 ± 26.10 Pa, while HA-Mal hydrogels with maleimide displayed a significantly higher compressive modulus of 152.3 ± 8.55 Pa (Fig. 2). After affibody conjugation, hydrogel compressive modulus did not differ significantly from that of the other hydrogels tested, indicating a negligible effect of the affibodies on HA hydrogel mechanical properties.

**Figure 2.**
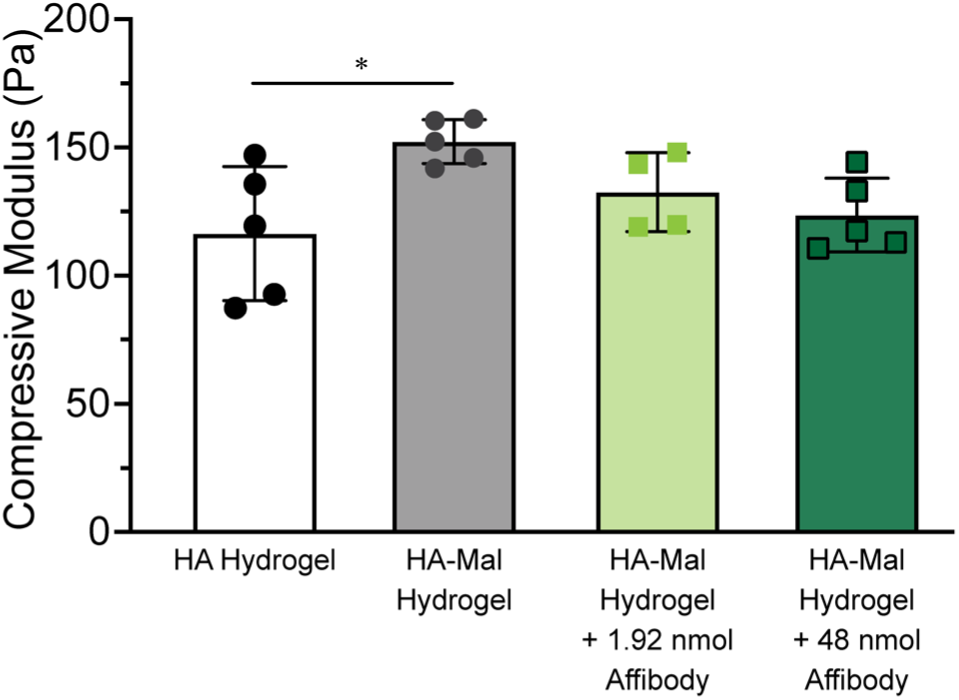
Compressive moduli of HA hydrogels with and without maleimide functional groups or affibodies. Formulations evaluated include HA hydrogels, HA-Mal hydrogels with maleimide-functionalized, and HA-Mal hydrogels with either 1.92 or 48 nmol of affibodies conjugated. Statistical significance between groups was determined using one-way ANOVA with Tukey’s post-hoc test. n=5-6 per group. *=p<0.05.

### 4.3 Collagen sponge reduces HA hydrogel swelling

The hydrogels and collagen sponge constructs were incubated in PBS at 37°C for 21 days, and their mass was measured periodically to evaluate swelling and/or degradation over time. Affibody-conjugated hydrogels swelled to 41.7% ± 3.6% above their original mass over 5 days before decreasing to 35.8% ± 4.3% above their original mass by day 21 (Fig. 3). Hydrogels without affibodies swelled maximally to 34.2% ± 2.9% above their original mass over 3 days and decreased to 23.4% ± 12.3% above their original mass by day 21. Collagen sponges lost 85.5% ± 2.5% of their hydrated initial mass after 1 day, remaining at 14.9% ± 1.1% of their original weight for the remainder of the experiment reflecting both loss of absorbed PBS from wicking and potentially some sponge degradation.

**Figure 3.**
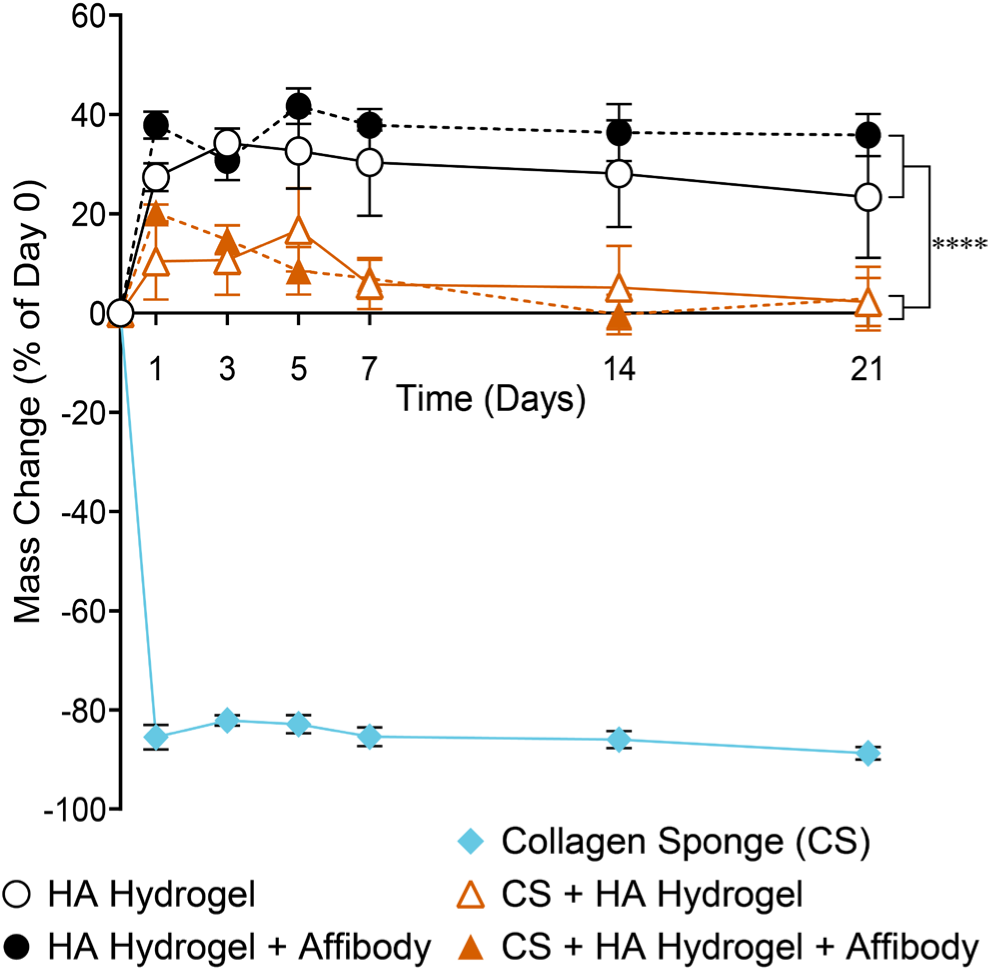
HA hydrogel and composite scaffold stability over 21 days. Mass change of HA hydrogels with and without affibodies, collagen sponges (CS), and composite HA hydrogels with and without affibodies formed within collagen sponges was measured for 21 days. Data are presented as mean ± standard deviation. Brackets indicate a significant main effect of the collagen sponge, comparing HA hydrogels with and without affibodies to composite scaffolds with and without affibodies, determined by three-way repeated-measures ANOVA with collagen sponge, affibody conjugation, and time as factors (Table S1). Collagen sponges were excluded from this analysis because they contain no HA hydrogel and would create an unbalanced three-way design. n=3 for the HA hydrogel group and n=4 for all other groups. ****=p<0.0001.

Forming the hydrogel within a collagen sponge significantly reduced construct swelling (p<0.0001) and was the largest contributor to differences in mass change among constructs. Affibody conjugation had no significant effect on mass change (p=0.10), and this held for hydrogels and composite scaffolds alike (collagen sponge × affibody interaction, p=0.14). Affibody-conjugated hydrogels crosslinked within collagen sponges swelled to 20.2% ± 1.7% above their original weight after 1 day and gradually returned to their original weight by day 21. Hydrogels without affibodies crosslinked within collagen sponges similarly swelled to 16.7% ± 8.4% above their original weight by day 5 and returned to their original weight by day 21. Overall, all composite scaffolds returned to their initial weight by day 21, while hydrogels without a collagen sponge remained swollen. Full statistical analysis results are provided in the supplemental information (Table S1).

### 4.4 Burst release of BMP-2 from collagen sponges was reduced by HA hydrogels

BMP-2 release from both HA hydrogels and composite collagen sponge + HA hydrogels were evaluated with and without affibody conjugation and compared to BMP-2 release from collagen sponges. 100 ng of BMP-2 were loaded onto the constructs and BMP-2 release was evaluated over 21 days. Initial encapsulation of BMP-2 was >94% for all groups regardless of the construct composition (Fig. 4A). Collagen sponges released 36.0% ± 1.2% of the loaded BMP-2 by day 21 (Fig. 4B). Conversely, HA hydrogels released 11.8% ± 0.7% of the loaded BMP-2 after 21 days. Affibody-conjugated HA hydrogels released less BMP-2 than collagen sponges and HA hydrogels, with 6.5% ± 0.9% being released by day 21. HA hydrogels with and without affibodies crosslinked within collagen sponges demonstrated further reduced cumulative BMP-2 release over 21 days of 4.9% ± 0.4% and 8.4% ± 1.8%, respectively. At day 21, collagen sponges had released more BMP-2 than every other construct (Tukey’s post-hoc, p<0.0001 for all comparisons). Affibody conjugation reduced cumulative release from HA hydrogels alone (Tukey’s post-hoc, p<0.001) but not in composite scaffolds (Tukey’s post-hoc, p=0.11).

**Figure 4.**
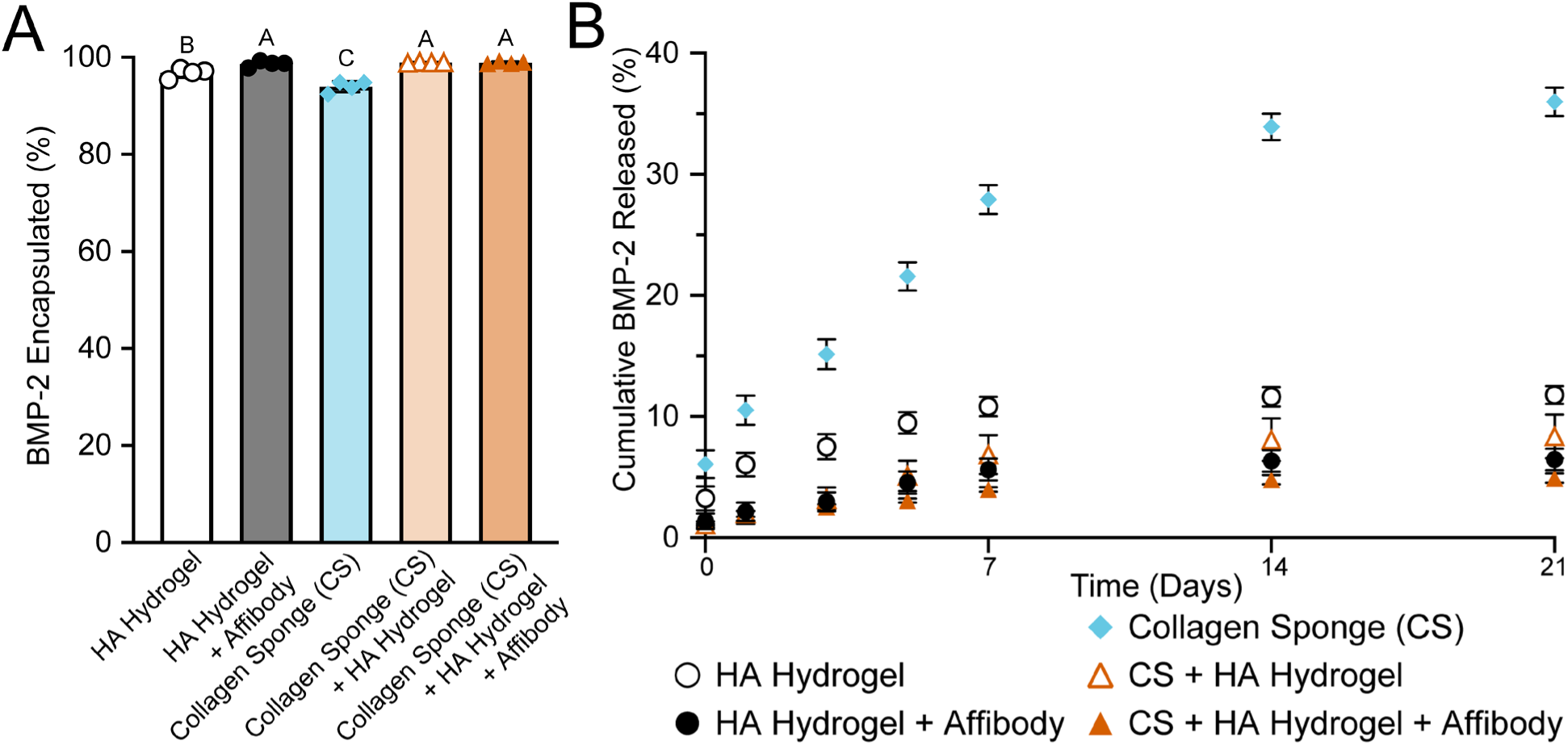
BMP-2 encapsulation and release from collagen sponges, HA hydrogels, and composite scaffolds with and without affibodies. A) Percent of BMP-2 encapsulated within HA hydrogels, affibody-conjugated HA hydrogels, collagen sponges (CS), CS + HA hydrogels, and CS + affibody-conjugated HA hydrogels. Groups that share a letter are not significantly different; groups with different letters are significantly different (p <0.05, one-way ANOVA with Tukey’s post-hoc test). B) Cumulative BMP-2 released from each construct over 21 days, expressed as a percentage of loaded BMP-2. Construct type, time, and the interaction between construct type and time all had significant effects on BMP-2 release (two-way repeated-measures ANOVA, p <0.0001, Table S2). Data are presented as mean ± standard deviation. n=4 per group.

### 4.5 Affibody-conjugated hydrogels with collagen sponges improved BMP-2 mediated subcutaneous mineralization

To evaluate how BMP-2-specific affibodies and BMP-2 dose influenced bone formation *in vivo*, we first evaluated ectopic BMP-2-mediated mineral formation in a subcutaneous implant model, which allowed us to screen several constructs and BMP-2 dose combinations at once and determine whether increased BMP-2 retention improved bone formation. Collagen sponges and hydrogels with and without BMP-2-specific affibodies formed within collagen sponges were loaded with either 2.5 µg or 5.0 µg of BMP-2 and implanted subcutaneously in the backs of female Sprague Dawley rats for 28 days. A 500-fold molar excess of affibodies to BMP-2 dose was conjugated to the HA-Mal-Ox, as this ratio of affibodies to BMP-2 has previously reduced BMP-2 release from HA and PEG-Mal hydrogels.[23,25] After four weeks, constructs were explanted, and bone formation was assessed using µCT (Fig. 5A). BMP-2-loaded collagen sponge implants exhibited bone volumes of 0.43 ± 0.65 mm³ with 2.5 µg of BMP-2 and 0.78 ± 0.68 mm³ with 5.0 µg of BMP-2 (Fig. 5B). Affibody-conjugated hydrogels within collagen sponges increased bone formation in a BMP-2 dose-dependent manner with average bone volumes of 2.20 ± 1.40 mm³ for 2.5 µg of BMP-2 and 6.45 ± 3.28 mm³ for 5.0 µg of BMP-2. 5.0 µg BMP-2-loaded composite scaffolds containing affibody conjugated HA hydrogels yielded significantly more bone than both BMP-2 doses delivered from collagen sponges and composite scaffolds without affibodies, suggesting that affibody-mediated BMP-2 retention enhances bone formation.

**Figure 5.**
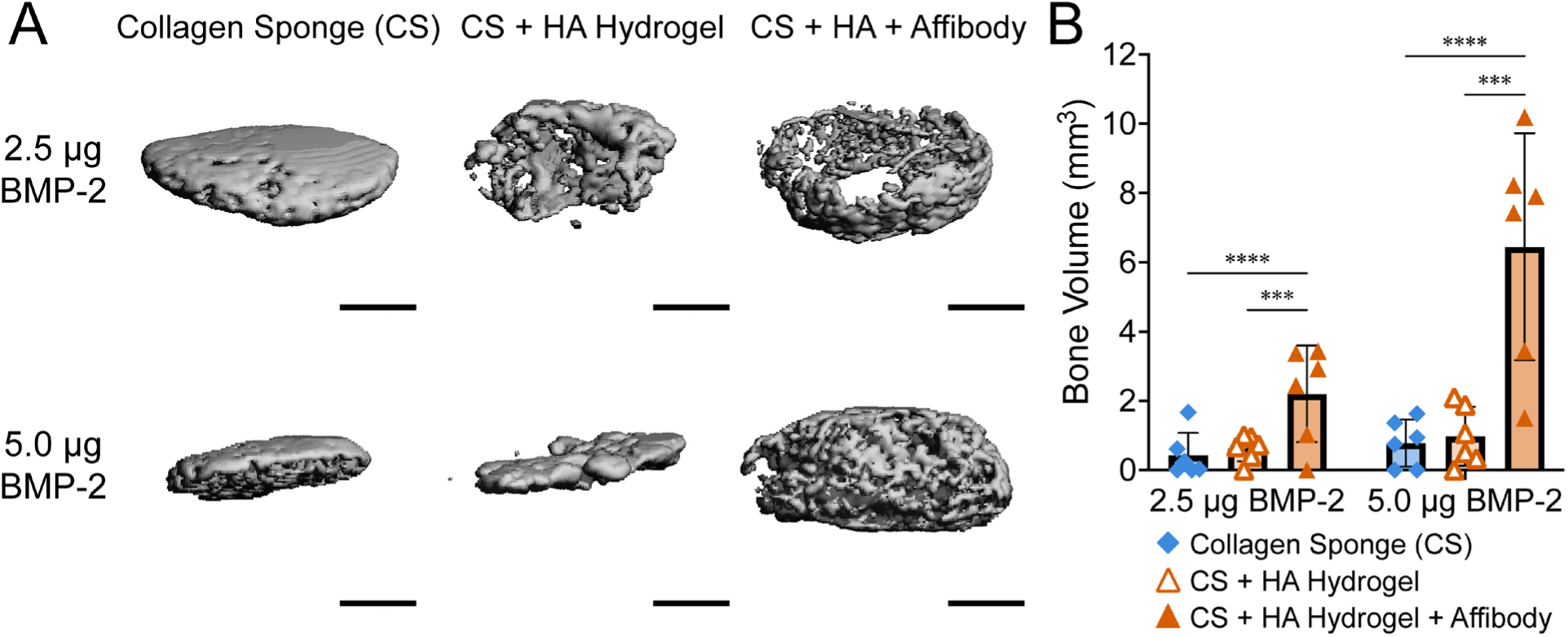
Affibody-conjugated HA hydrogels enhance ectopic mineralization in a BMP-2 dose-dependent manner. Collagen sponges (CS), CS + HA hydrogels, and CS + affibody-conjugated HA hydrogels were loaded with 2.5 or 5.0 µg of BMP-2 and implanted subcutaneously for 4 weeks. A) Representative µCT reconstructions of explanted constructs depicting mineralization. Scale bar is 1 mm. B) Bone volume quantified by µCT. Statistical significance was determined using two-way ANOVA with Tukey’s post-hoc test. *** = p <0.001, **** = p <0.0001. Data are presented as mean ± standard deviation. n=6 per group.

### 4.6 Histological evaluation of subcutaneous implants

Representative samples based on the mean bone volume from each group were selected for qualitative histological analysis. Hematoxylin and eosin-y (H&E) staining of subcutaneous implants after 28 days revealed similar disk-like morphology and cellular infiltration for all constructs regardless of BMP-2 dose (Fig. S5). However, enlarged views of the implant interiors revealed differences in the spatial distribution of cells and structures among the constructs (Fig. 6). At both BMP-2 doses, collagen sponge implants had porous interiors with dense matrix deposition (black arrows) primarily around the implant periphery (Fig. 6A, 6D). In contrast, the incorporation of HA hydrogels within the collagen sponge resulted in more extensive cellular infiltration towards the interior of the implants (white arrows). Notably, collagen sponges with affibody-conjugated HA hydrogels containing 5.0 µg of BMP-2 displayed a trabecular-like network (asterisks) towards the center of the implants (Fig. 6F). This trabecular-like morphology suggests that affibody-mediated BMP-2 retention may support more organized bone formation within the implant interior rather than the predominantly peripheral mineralization observed with the collagen sponge implants. These observations are consistent with the higher bone volume measured by µCT in affibody-conjugated constructs (Fig. 5B).

**Figure 6.**
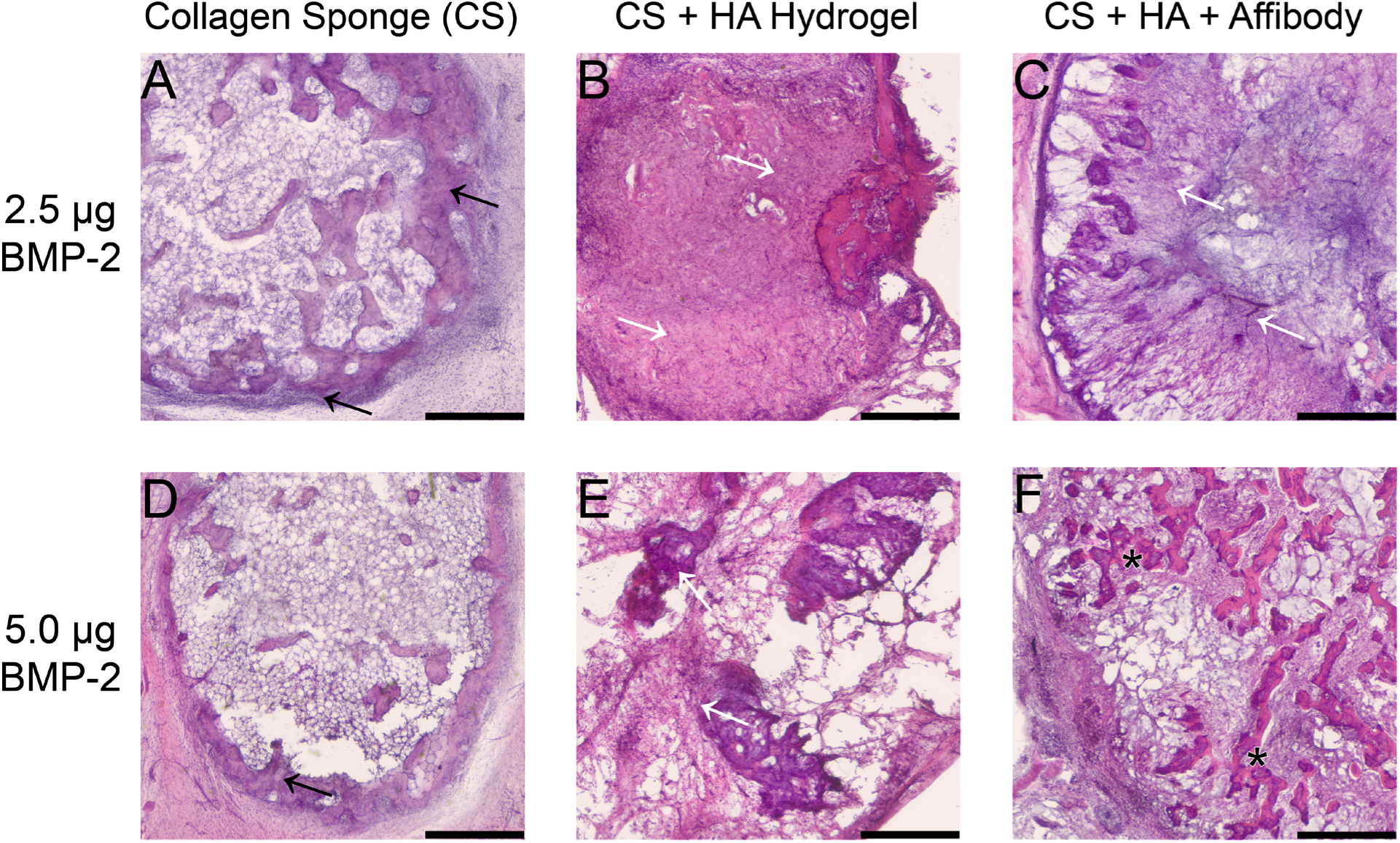
Representative histology from subcutaneous implants. 2.5 µg of BMP-2 was loaded into A) collagen sponge, B) collagen sponge and HA hydrogel, and C) collagen sponge and affibody-conjugated HA hydrogel. 5.0 µg of BMP-2 was also evaluated in D) collagen sponge, E) collagen sponge and HA hydrogel, and F) collagen sponge and affibody-conjugated HA hydrogel. Black arrows indicate matrix deposition at the implant periphery in collagen sponge only groups while white arrows indicate cellular infiltration toward the implant interior seen in composite scaffolds. Asterisks indicate trabecular-like structures. Scale bar is 500 µm.

### 4.7 BMP-2-loaded composite scaffolds improve bone healing in critically sized femoral defects in rats

We implanted collagen sponges, composite scaffolds containing collagen sponges and HA hydrogels without affibodies, and composite scaffolds containing collagen sponges and HA hydrogels conjugated with BMP-2-specific affibodies in 6-mm critically sized femoral defects in Sprague Dawley rats. All constructs were loaded with 1.0 µg of BMP-2, which has resulted in bone formation with variable bone bridging in this model.[13] Serial radiographs taken at 4, 8, and 12 weeks post-injury showed progressive mineralization in all treatment groups (Fig. 7). Based on 12-week radiographs, bone bridging was observed in all defects treated with BMP-2-loaded composite scaffolds (11 of 11) compared with 2 of 4 defects treated with BMP-2-loaded collagen sponges (Fisher’s exact test, p=0.057).

**Figure 7.**
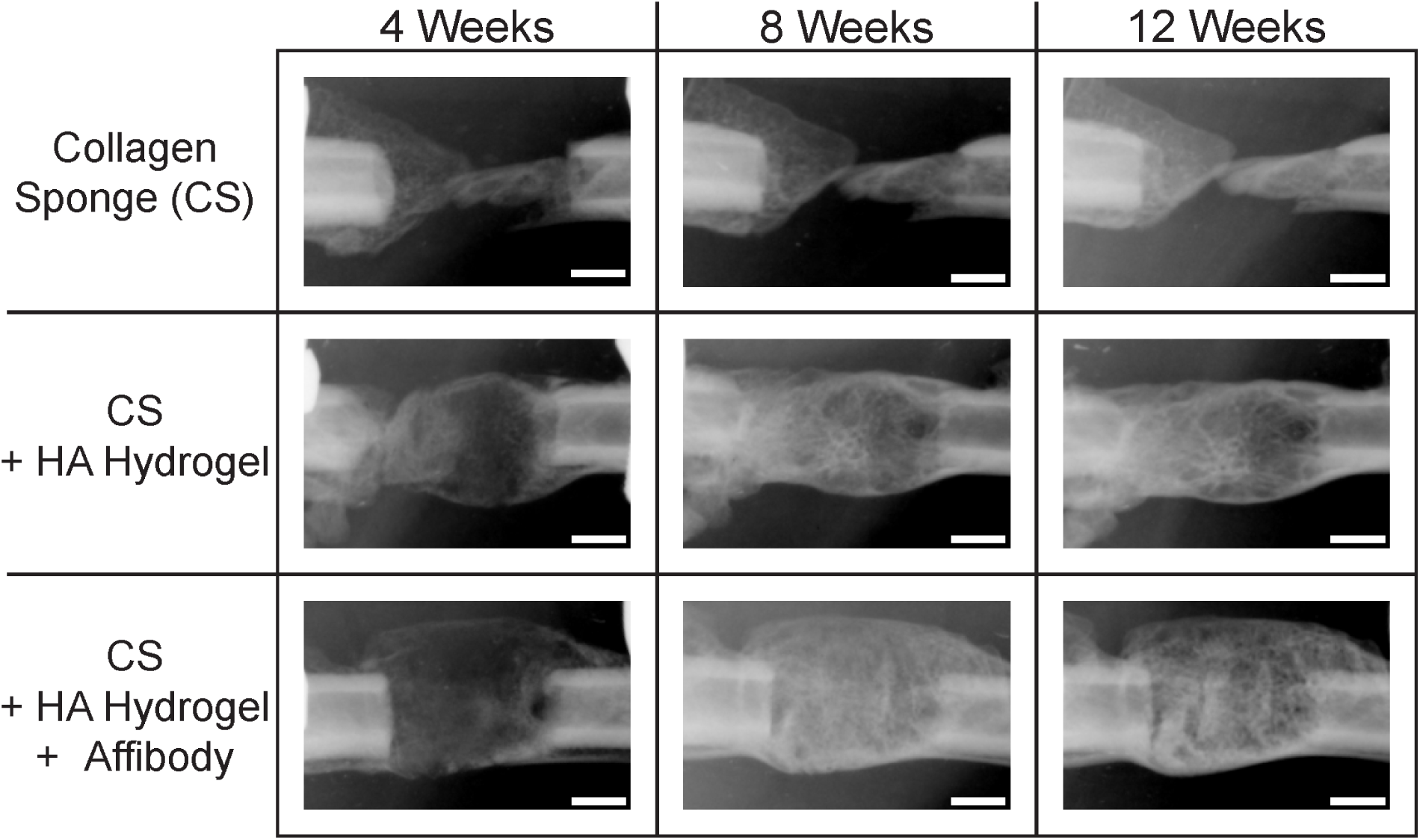
Representative serial radiographs of critically sized 6-mm femoral defects treated with 1.0 µg BMP-2-loaded collagen sponges and composite scaffolds containing collagen sponges and HA hydrogels with and without affibodies. Scale bar is 2 mm.

µCT was performed *in vivo* at 6 weeks and *ex vivo* at 12 weeks to quantify total bone volume in the defect space. At 6 weeks, femoral defects treated with affibody-conjugated composite scaffolds exhibited significantly higher bone volume than those treated with collagen sponges (Fig. 8A). At 12 weeks, both composite scaffolds with and without affibodies formed more mineral than collagen sponges (Fig. 8E). Representative mid-sagittal sliced µCT reconstructions of femoral defects treated with collagen sponges (Fig. 8B), composite collagen sponge and HA hydrogels (Fig. 8C), and composite scaffolds with affibody-conjugated HA hydrogels (Fig. 8D) at 6 weeks were consistent with radiographic evidence of bone growth throughout the defect space. Representative mid-sagittal sliced µCT reconstructions of the same femurs at 12 weeks (Fig. 8F-H) depicted increased defect bone volume in groups treated with composite scaffolds with and without affibodies compared to collagen sponge alone.

**Figure 8.**
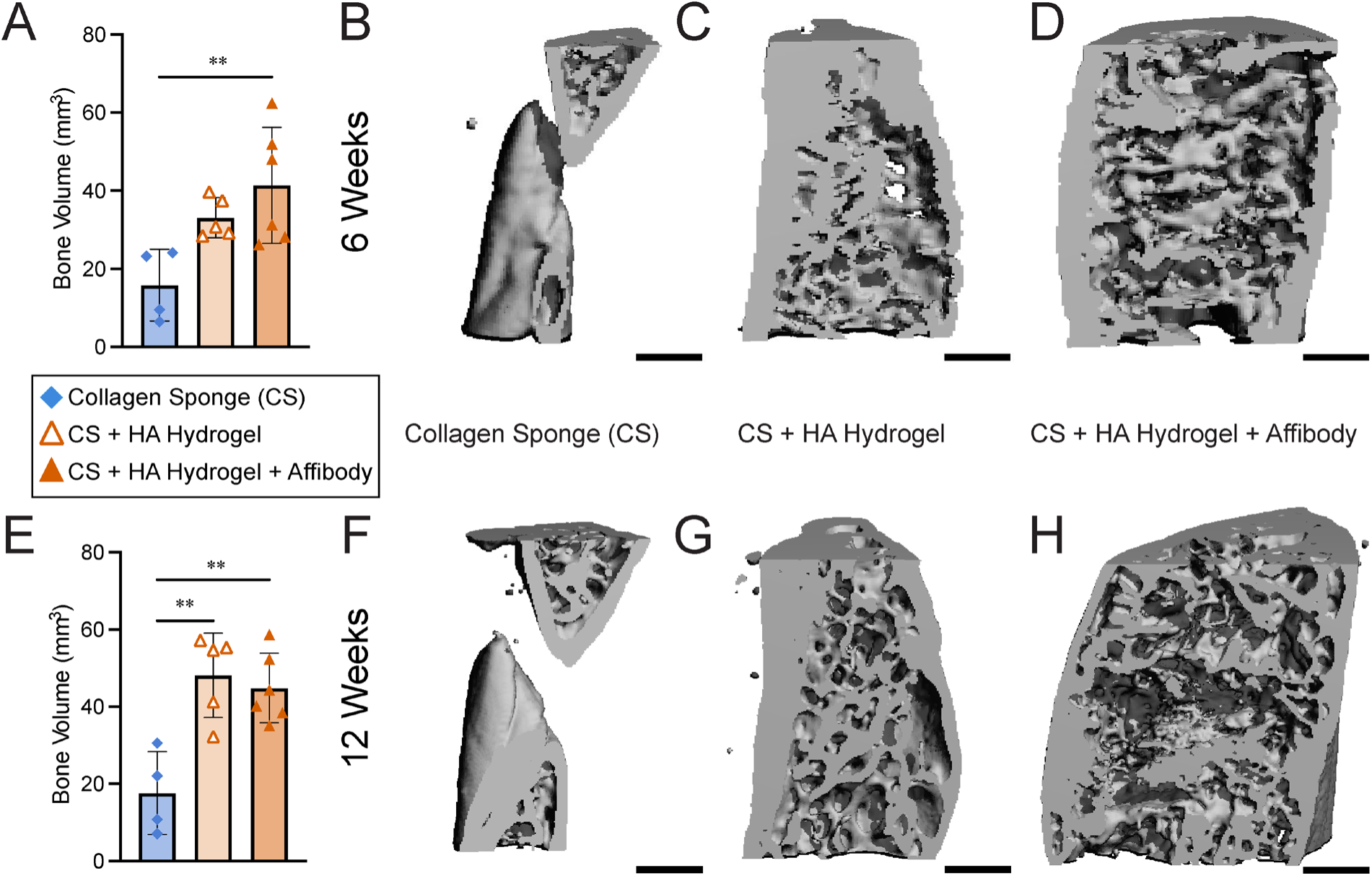
µCT evaluation of regenerated bone in a femoral defect model at 6 and 12 weeks post-injury. Critically sized 6-mm defects were treated with 1.0 µg of BMP-2 using collagen sponges, composite collagen sponge and HA hydrogels, and composite scaffolds with affibody-conjugated HA hydrogels. A) *In vivo* µCT quantification of total new bone volume formed after 6 weeks. Representative mid-sagittal sliced µCT reconstructions for B) collagen sponge, C) collagen sponge and HA hydrogels, and D) collagen sponge and affibody-conjugated HA hydrogels after 6 weeks. E) *Ex vivo* µCT quantification of total new bone volume formed after 12 weeks. Representative mid-sagittal sliced µCT reconstructions for F) collagen sponge, G) collagen sponge and HA hydrogels, and H) collagen sponge and affibody-conjugated HA hydrogels after 12 weeks. Statistical significance between groups was determined using a one-way ANOVA with Tukey’s post-hoc test. ** = p <0.01. n=4-6 per group. Scale bar is 1 mm.

### 4.8 Histological evaluation of femoral bone defects

Histological staining was performed on the same representative femurs used for µCT reconstructions. Hematoxylin and eosin (H&E) staining revealed more robust tissue formation in defects treated with HA hydrogels formed within collagen sponges with and without affibodies compared to collagen sponges alone (Fig. 9A-C). Adjacent sections were stained with Picrosirius Red and imaged under polarized light to visualize collagen, revealing similar types and amounts of collagen in each treatment group (Fig. 9D-I), with green birefringence indicating thinner less organized collagen fibers and red/yellow birefringence indicating thicker more mature fibers.[33] Samples stained with Safranin O and Fast Green revealed no proteoglycan staining (Fig. S6), which was expected and consistent with our previous work; by 12 weeks, cartilage produced at earlier stages of bone repair would have been remodeled.[24,34,35]

**Figure 9.**
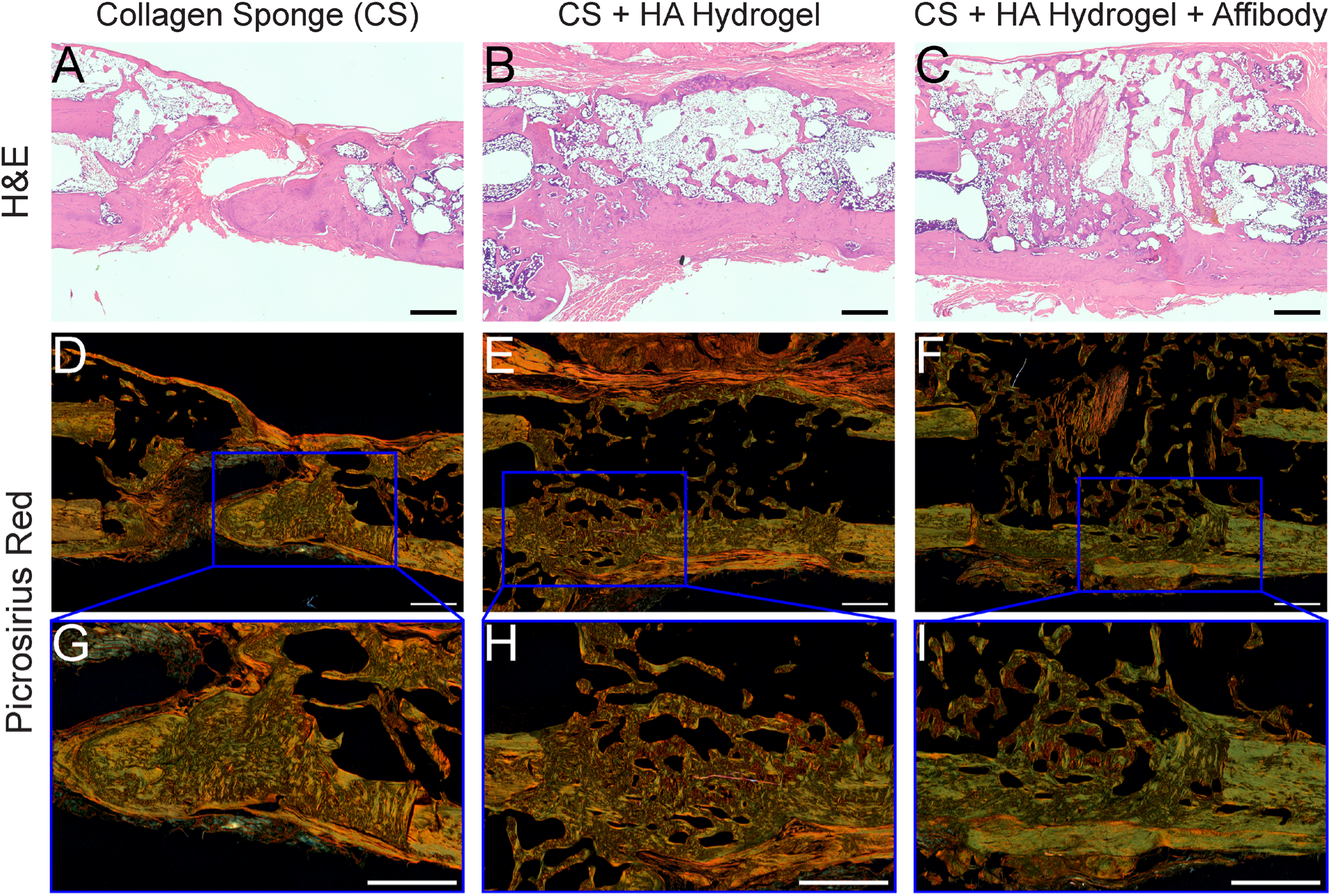
Histological staining of femoral bone defects at 12 weeks post-injury. Critically sized 6-mm defects were treated with 1.0 µg of BMP-2 loaded into collagen sponges, composite collagen sponge and HA hydrogels, and composite scaffolds with affibody-conjugated HA hydrogels. A-C) Hematoxylin and Eosin (H&E) staining of treated femurs, imaged with a 5x objective under brightfield. D-F) Picrosirius Red staining of adjacent sections, imaged with a 10x objective under polarized light. G-I) Enlarged views of the regions outlined in blue in D-F, near the intact bone ends of the defects. Scale bar is 1 mm.

## 5. Discussion

Despite inconsistent clinical evidence from spinal fusion procedures, BMP-2 remains widely used because it is osteoinductive, reduces surgical time, and avoids the donor site complications associated with autograft harvest.[36,37] However, the collagen sponge used as a BMP-2 carrier retains BMP-2 through weak electrostatic interactions, resulting in burst release of the protein, and the compensatory supraphysiological doses used are associated with soft tissue inflammation, heterotopic ossification, and other implant-site complications.[6] The criteria for an improved carrier for BMP-2-mediated bone regeneration include the ability to better localize protein within the implant site, allow cell and tissue infiltration and remodeling throughout the bone healing process, and be compatible with the surgical workflow established in the clinic. Here, we developed a hydrazone crosslinked HA hydrogel that can be formed *in situ* within clinically used collagen sponges and includes a maleimide functional group for conjugation of BMP-2-specific affibodies to improve BMP-2 retention within the biomaterial. Maleimide functionalization modestly increased the compressive modulus of the hydrogel. Furthermore, subsequent affibody conjugation had no impact on compressive modulus, indicating that neither functionalization nor conjugation compromised hydrazone crosslinking. Forming the hydrogel within the collagen sponge significantly reduced construct swelling, which reduces protein burst release and hydrolytic hydrogel degradation. *In vitro,* composite scaffolds retained substantially more BMP-2 than the collagen sponge alone, and conjugation of BMP-2-specific affibodies further reduced cumulative BMP-2 release. *In vivo,* affibody conjugation improved BMP-2-mediated ectopic bone formation in subcutaneous implants. In a critically sized rat femoral defect, BMP-2-loaded composite scaffolds with and without affibodies both significantly increased bone volume relative to BMP-2-loaded collagen sponges at 12 weeks, and all defects treated with composite scaffolds bridged.

Localizing BMP-2 within the construct through affibody-mediated affinity interactions requires conjugating enough affibodies to the hydrogel to bind and retain the BMP-2 dose necessary to stimulate bone formation. This has been a challenge with other hydrogels, especially when the same functional groups are used for both affibody conjugation and hydrogel crosslinking.[25,38,39] Therefore, a critical objective of this work was to decouple affibody conjugation from hydrogel crosslinking. We previously attempted affibody conjugation using photoinitiated thiol-norbornene chemistry; however, poor conjugation efficiency, limited polymer solubility, and inconsistent network formation prevented delivery of the BMP-2 doses required for *in vivo* bone regeneration. In this work, we conjugated BMP-2-specific affibodies to HA via thiol-Michael addition between the C-terminal cysteine on the affibody and the maleimide on the HA polymer without affecting hydrogel crosslinking or requiring UV light and a photoinitiator. This conjugation strategy allowed us to maintain a 500-fold molar excess of affibodies to BMP-2 as the dose increased from 0.1 µg previously delivered *in vitro* [25] to 1.0-5.0 µg required *in vivo*, representing a 50-fold increase in affibody conjugation.

Other maleimide-functionalized HA hydrogels have also been developed for *in vivo* bone regeneration, where the maleimide group was used to crosslink the hydrogel.[40] However, these hydrogels required the conjugation of matrix metalloproteinase (MMP)-sensitive crosslinkers and cell-adhesive peptides to promote cell infiltration and material degradation [14,28], since biomaterial carriers for delivering BMP-2 must allow cell and tissue infiltration to stimulate bone formation and remodeling.[41] In contrast, the hydrazone crosslinking used here is orthogonal to the thiol-maleimide reaction, leaving the maleimide groups available for affibody conjugation, and involves dynamic covalent bonds that can be easily remodeled by cells.[42] Furthermore, we formed the HA hydrogel within the absorbable collagen sponge, which allows cell and tissue infiltration and is already clinically approved for BMP-2 delivery. The resulting composite scaffolds were easier to handle during surgical implantation than HA hydrogels and swelled significantly less. Swelling changes hydrogel mesh size, which impacts both protein diffusion and cellular infiltration, and excess swelling can negatively impact surrounding cells and tissues.[14–17,43,44] Affibody-conjugated hydrogels swelled to 41.7% ± 3.6% above their initial mass by day 5 and decreased to 35.8% ± 4.3% above their original mass by day 21. In contrast, the same hydrogels formed within a collagen sponge swelled by 20.2% ± 1.7% of their initial mass and gradually returned to their initial mass over 21 days. Collagen sponges lost 85.5% ± 2.5% of their hydrated initial mass after 1 day. Thus, forming the HA hydrogels within collagen sponges constrained initial swelling and preserved the collagen sponges over 21 days.

*In vitro* BMP-2 release from affibody-conjugated hydrogels was lower than BMP-2 release from sponges and hydrogels without affibodies. Over 21 days, collagen sponges released 36.0% ± 1.2% of their loaded BMP-2, compared to 11.8% ± 0.7% from HA hydrogels, 8.4% ± 1.8% from composite scaffolds, and 4.9% ± 0.4% from affibody-conjugated composite scaffolds. While our previous study demonstrated that affibodies increased protein retention *in vivo*, this is the first study to demonstrate that affibody-mediated BMP-2 delivery improves bone formation.[24] Affibody-conjugated composite scaffolds produced the highest bone volumes ectopically among groups at both 2.5 µg and 5.0 µg of BMP-2. Affibody-conjugated composite scaffolds were also the only group in which mineralization increased significantly with increasing BMP-2 dose, suggesting that affibody conjugation, rather than the hydrogel matrix alone, drove the dose-dependent increase in ectopic bone formation.

We next evaluated BMP-2-loaded collagen sponges and composite scaffolds with and without affibodies in a critically sized rat femoral defect, which is a well-established model for evaluating bone regeneration therapies.[19,45–47] Both composite scaffolds outperformed the collagen sponge at 12 weeks, and the two did not differ from each other at either 6 or 12 weeks. All defects treated with composite scaffolds bridged, compared with half of those treated with collagen sponges, though this did not reach significance (p=0.057), reflecting the small number of animals remaining in the collagen sponge group after two exclusions for fixation plate failure. The improvement of both composite scaffolds over the collagen sponge may be attributed to the presence of the HA network, which slowed BMP-2 release *in vitro*. Affibody-conjugated composite scaffolds were the only group to form significantly more bone than the collagen sponge at 6 weeks, which is consistent with affinity binding increasing BMP-2 retention. Affibody-mediated BMP-2 retention would be expected to have the largest effect on bone repair at the early stages of healing while exogenous BMP-2 is present and driving cell recruitment and differentiation during the fibrocartilaginous callus formation stage.[48] However, because bone formation induced by the two composite scaffolds did not differ significantly at 6 weeks, the early benefit of affibody conjugation in this model remains unconfirmed. By 12 weeks, composite scaffolds without affibodies reached the same bone volume as affibody-conjugated composite scaffolds. A similar effect occurred in our previous work, in which hydrogels delivering 5.0 µg of BMP-2 bridged every critically sized femoral defect regardless of affibody conjugation.[24]

While these results demonstrate that composite scaffolds improve BMP-2-mediated bone regeneration relative to the collagen sponge, the contribution of affibody conjugation *in vivo* must be further investigated. Histology and immunohistochemistry performed at additional time points during the early stages of bone healing (4-6 weeks) could provide valuable information about early effects of prolonging local BMP-2 retention via affinity interactions. Early callus formation can be evaluated by staining for osteoprogenitor cells, vascular ingrowth, and macrophage infiltration. Future studies should test a lower BMP-2 dose to assess whether affibody conjugation improves bone formation at doses where the composite construct alone is no longer sufficient for bridging a critically sized femoral defect. At 1.0 µg of BMP-2, every defect treated with a composite scaffold bridged regardless of affibody conjugation, which could not resolve the benefits of affibody conjugation from improved retention provided from the HA hydrogel. At a dose where the composite scaffold alone bridges only some defects, differences between affinity-mediated BMP-2 delivery and retention could be disentangled.

## 6. Conclusion

In this work, we developed a novel maleimide functionalized, hydrazone crosslinked hyaluronic acid hydrogel that employs orthogonal reactions for affibody conjugation and network crosslinking. Conjugating BMP-2-specific affibodies using a thiol-Michael addition reaction required no photoinitiator or ultraviolet light and did not compromise hydrogel properties. Forming the hydrogel within a clinically used collagen sponge produced a composite construct that could be press-fit into the defect, swelled less than the hydrogel alone, and retained more BMP-2 *in vitro* than the collagen sponge. In a subcutaneous ectopic implant model, affibody conjugation improved BMP-2-mediated bone formation at both doses tested, demonstrating for the first time that conjugating BMP-2-specific affibodies to a biomaterial delivery vehicle improves bone formation. In a critically sized femoral defect treated with 1.0 µg of BMP-2, composite scaffolds with and without affibodies both increased bone volume and defect bridging relative to the collagen sponge, although the two composite scaffolds did not differ from each other. Together, these results establish a natural polymer hydrogel platform for affinity-based BMP-2 delivery that builds on, rather than replaces, the collagen sponge used clinically. Because affibody density and crosslinking are controlled by orthogonal chemistries, the same platform can be used with additional affinity binders to deliver other growth factors.

## 7. Acknowledgments and Funding

We are grateful for funding from the National Institutes of Health (R35 Maximizing Investigators’ Research Award (MIRA), R35-GM147507 to M.H.H.), Department of Defense (Discovery Award, Peer-Reviewed Medical Research Program W81XWH2210700 to M.H.H.), and the Lary Simpson Professorship to M.H.H. We are also grateful for funding from the Wu Tsai Human Performance Alliance and the Joe and Clara Tsai Foundation to M.H.H. and N.W. Y.C.P. was supported by the National Science Foundation Graduate Research Fellowship Program (NSF GRFP). We thank members of the Hettiaratchi and Willett labs for their assistance with animal surgeries and feedback on this manuscript. We also thank Dr. Danielle Benoit for the use of her lab’s peptide synthesizer and HPLC system.

