## Supplemental Information for "Affibody-conjugated hyaluronic acid hydrogels within collagen sponges enhance dose-dependent BMP-2-mediated bone formation and repair of critically sized femoral defects"

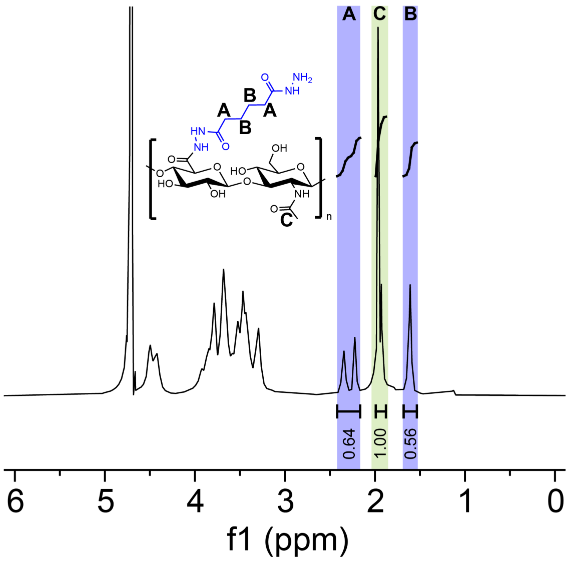


Figure S1. Representative $^{1}$H-NMR spectrum of HA-Mal depicting modification of HA with maleimide functional groups.

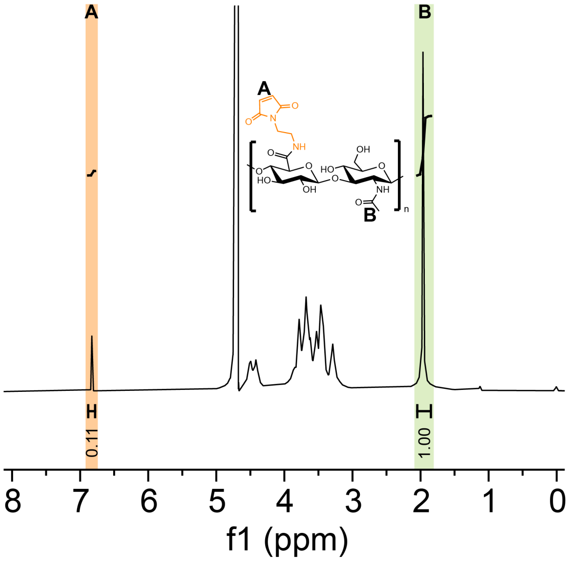


Figure S2. Representative $^{1}$H-NMR spectrum of HA-ADH depicting modification of HA with adipic acid dihydrazide functional groups.

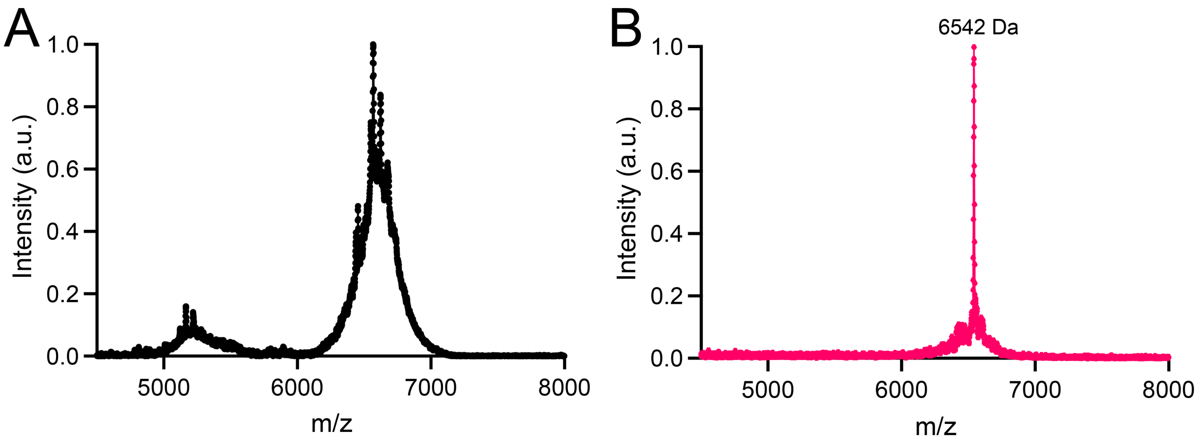


Figure S3. Representative MALDI-TOF spectra of crude and pure BMP-2-specific affibodies. A) Crude affibodies after cleavage from Fmoc resin. B) Affibodies purified using HPLC with expected mass at 6542 Da.

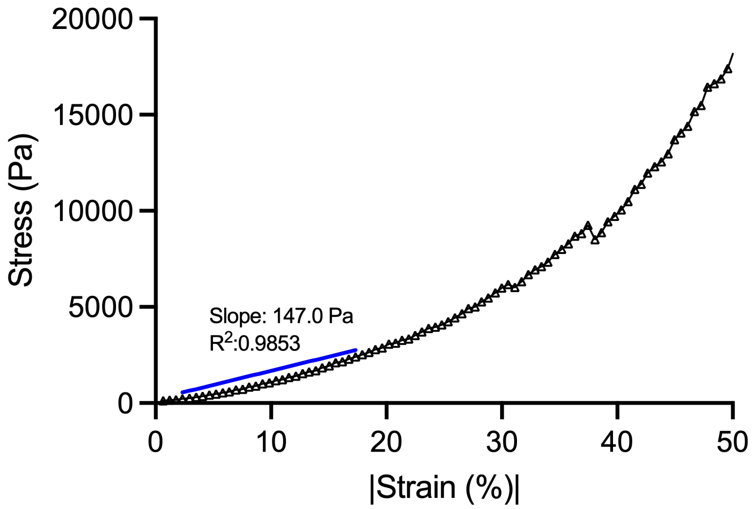


Figure S4. Representative stress-strain curve of compression of HA hydrogel. Representative stress vs. strain curve of HA hydrogels. The slope of the curve in the linear region at 15% strain was used to calculate compressive modulus for each hydrogel formulation.

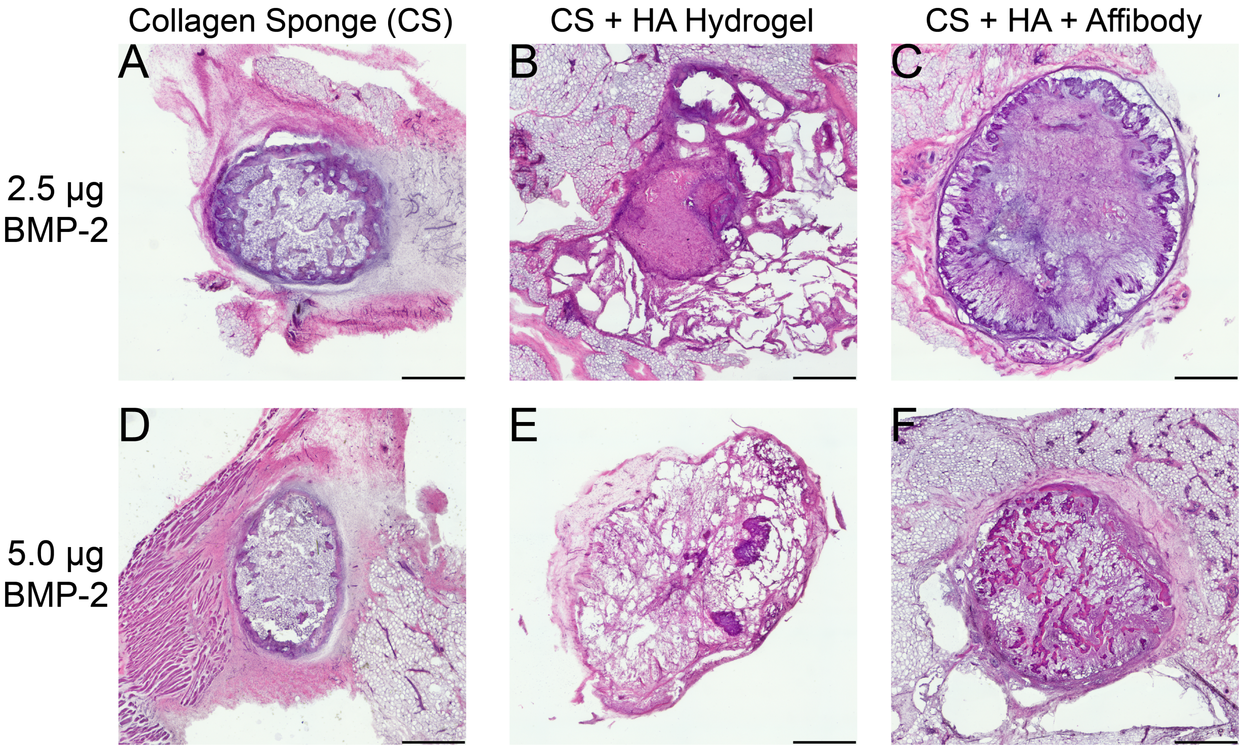


Figure S5. Representative histology from subcutaneous implants. 2.5 µg of BMP-2 was loaded into A) collagen sponge, B) collagen sponge and HA hydrogel, and C) collagen sponge and affibody-conjugated HA hydrogel. 5.0 µg of BMP-2 was also evaluated in D) collagen sponge, E) collagen sponge and HA hydrogel, and F) collagen sponge and affibody-conjugated HA hydrogel. All constructs showed similar disk-like morphology and cellular infiltration after 28 days. Scale bar is 1 mm.

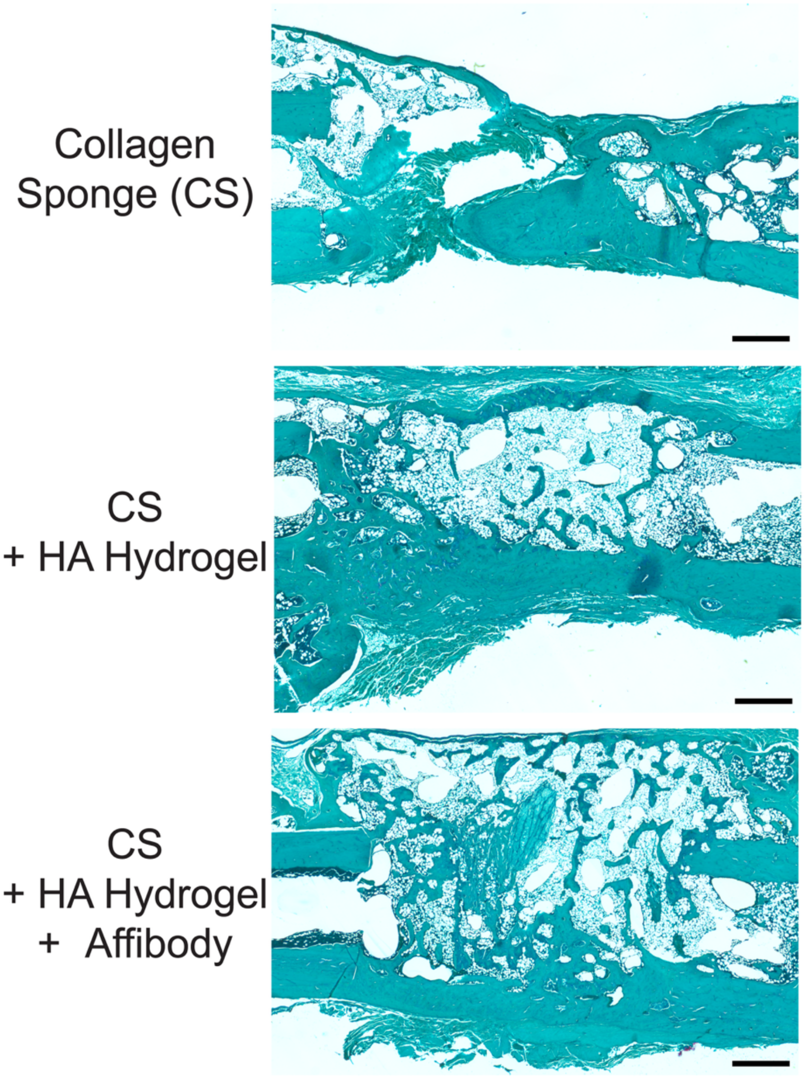


Figure S6. Safranin O staining of femoral defects at 12 weeks post-injury. Defects were treated with 1.0 μg of BMP-2 using collagen sponges, composite collagen sponge and HA hydrogels, and composite scaffolds with affibody-conjugated HA hydrogels. Scale bar is 1mm.

| Source of variation | % of total variation | F ($DF_n$, $DF_d$) | p value |
|---|---|---|---|
| Collagen sponge | 69.20 | F(1, 11) = 124.7 | < 0.0001 |
| Affibody conjugation | 1.74 | F(1, 11) = 3.136 | 0.1042 |
| Collagen sponge × affibody conjugation | 1.44 | F(1, 11) = 2.589 | 0.1359 |
| Time | 5.03 | F(3.346, 36.80) = 8.510 | 0.0001 |
| Time × collagen sponge | 2.31 | F(3.346, 36.80) = 3.913 | 0.0133 |
| Time × affibody conjugation | 1.51 | F(3.346, 36.80) = 2.556 | 0.0643 |
| Time × collagen sponge × affibody conjugation | 2.07 | F(3.346, 36.80) = 3.507 | 0.021 |

Table S1. Three-way repeated-measures ANOVA of construct mass change over 21 days. Factors were collagen sponge (present or absent), affibody conjugation (present or absent), and time (days 1, 3, 5, 7, 14, and 21; repeated measure). Collagen sponges alone were excluded from the analysis. n = 3-4 per group. P value was calculated from $F$ statistic with $DF_n$ and $DF_d$ denoting the degrees of freedom in the numerator and denominator respectively.

| Source of variation | % of total variation | F ($DF_n$, $DF_d$) | p value |
|---|---|---|---|
| Construct | 62.19 | F(4, 15) = 271.9 | < 0.0001 |
| Time | 20.67 | F(1.219, 18.29) = 3313 | < 0.0001 |
| Time × construct | 16.19 | F(4.878, 18.29) = 648.5 | < 0.0001 |

Table S2. Two-way repeated-measures ANOVA of cumulative BMP-2 release over 21 days. Factors were construct type (HA hydrogels, affibody-conjugated HA hydrogels, collagen sponges, and composite scaffolds with and without affibodies) and time (days 0, 1, 3, 5, 7, 14, and 21; repeated measure). n = 4 per group. P value was calculated from $F$ statistic with $DF_n$ and $DF_d$ denoting the degrees of freedom in the numerator and denominator respectively.
